# Oxidative Stress and Neuroinflammation in a Rat Model of Co-Morbid Obesity and Psychogenic Stress

**DOI:** 10.1101/2020.05.19.104794

**Authors:** Jose M. Santiago Santana, Julio David Vega-Torres, Perla Ontiveros Angel, Jeong Bin Lee, Yaria Arroyo Torres, Alondra Y. Cruz Gonzalez, Esther Aponte Boria, Deisha Zabala Ortiz, Carolina Alvarez Carmona, Johnny D. Figueroa

## Abstract

**Background:** There is growing recognition for a reciprocal, bidirectional link between anxiety disorders and obesity. Although the mechanisms linking obesity and anxiety remain speculative, this bidirectionality suggests shared pathophysiological processes. Neuroinflammation and oxidative damage are implicated in both pathological anxiety and obesity. This study investigates the relative contribution of comorbid diet-induced obesity and stress-induced anxiety to neuroinflammation and oxidative stress.

**Methods:** Thirty-six (36) male Lewis rats were divided into four groups based on diet type and stress exposure: 1) control diet unexposed (CDU) and 2) exposed (CDE), 3) Western-like high-saturated fat diet unexposed (WDU) and 4) exposed (WDE). Neurobehavioral tests were performed to assess anxiety-like behaviors. The catalytic concentrations of glutathione peroxidase and reductase were measured from plasma samples, and neuroinflammatory/oxidative stress biomarkers were measured from brain samples using Western blot. Correlations between behavioral phenotypes and biomarkers were assessed with Pearson’s correlation procedures.

**Results:** We found that WDE rats exhibited markedly increased levels of glial fibrillary acidic protein (185%), catalase protein (215%), and glutathione reductase (GSR) enzymatic activity (418%) relative to CDU rats. Interestingly, the brain protein levels of glutathione peroxidase (GPx) and catalase were positively associated with body weight and behavioral indices of anxiety.

**Conclusions:** Together, our results support a role for neuroinflammation and oxidative stress in heightened emotional reactivity to obesogenic environments and psychogenic stress. Uncovering adaptive responses to obesogenic environments characterized by high access to high-saturated fat/high-sugar diets and toxic stress has the potential to strongly impact how we treat psychiatric disorders in at-risk populations.

**Highlights:**

- Predatory odor stress heightens footshock reactivity and anxiety-like behaviors in Lewis rats.
- WD intake increases glutathione reductase activity in plasma.
- WD intake and PS exposure acted synergistically to increase the brain protein levels of catalase and the glial fibrillary acidic protein.
- The protein levels and activities of some redox/neuroinflammatory biomarkers are closely associated with behavioral proxies related to fear and anxiety in rats.

## Introduction

The prevalence of trauma exposure during adolescence is concerning and has severe consequences for the emergence of psychiatric disorders [1]. While the majority of traumatized children are not chronically affected, it is estimated that approximately one-third of trauma-exposed adolescents will develop anxiety symptomatology and posttraumatic stress disorder (PTSD) [2,3]. A better understanding of specific vulnerability factors may facilitate the development of tailored preventive and curative interventions. Obesity and the consumption of imbalanced diets rich in saturated fats and sugars have emerged as risk factors for the development of anxiety and stress-related disorders [4–11]. We have modeled this relationship by measuring stress reactivity to traumatic stress in a rat model of diet-induced obesity (DIO). Consistent with studies in obese adolescents, we reported that adolescent diet-induced obesity (DIO) in rats: **1)** reduces hippocampal volume [12], **2)** impairs the maturation of the corticolimbic fear circuits [13], **3)** enhances behavioral vulnerabilities to psychosocial stress [12–14], and **4)** results in profound fear extinction learning deficits [13,14], even in the absence of an obesogenic phenotype [15]. However, the molecular factors underlying these vulnerabilities are poorly understood.

Several lines of evidence indicate that obesity and the consumption of obesogenic diets lead to brain inflammation [16–18]. We demonstrated that the intake of an obesogenic diet upregulates the expression of the glial fibrillary acidic protein (GFAP) in the brain [13]. GFAP expression has been associated with activated astrocytes, which can lead to astrogliosis. Although the implications of this (mal)adaptive astrocytic response to obesogenic diets and brain insults remain controversial [19–23], astrocytes play critical roles in regulating oxidative stress in the brain [24,25]. Unlike neurons, astrocytes use glycolysis for energy production and, as a consequence, have been associated with a higher generation of mitochondrial reactive oxygen species (ROS) [26]. It is known that ROS accumulate following exposure to obesogenic diets and obesity in rodents and humans [27–31]. Due to its high lipid content, the brain is vulnerable to ROS damage, which can have devastating effects on brain function and in the etiology of anxiety and stress-related disorders [32–35]. Together, these findings underscore the importance of oxidative stress as a potential molecular link connecting obesity to increased stress vulnerability.

The functional coupling between astrocytes and neurons contributes to redox metabolism, which is tightly regulated by antioxidant systems [26]. Influential members of this defense system include catalase (CAT), glutathione peroxidase (GPx), glutathione reductase (GSR), and superoxide dismutases (SOD) [36,37]. In general, the antioxidant defense system works as follows: SOD converts O_2_. into hydrogen peroxide (H_2_O_2_), which is transformed into water by CAT and GPx [38,39]. Otherwise, H_2_O_2_ can react with iron and produce OH. in a process known as the Fenton reaction. GSR’s role relies on regenerating the reduced glutathione product of the GPx following GPx activities on lipid and non-lipid hydro-peroxides [40]. Studies demonstrate that lack of activity on this system leads to ROS buildup, inflammation, and may represent an important predisposing factor for anxiety and stress-related neuropsychiatric disorders [32–35].

Given that obesogenic diets induce oxidative stress, it is logical to hypothesize that the antioxidant network will mount a compensatory response to preserve redox homeostasis. Therefore, examining the response of the antioxidant system to obesogenic diets may provide valuable insight into the potential contribution of oxidative stress to anxiety and other stress-related psychopathologies. This study is the first to investigate the combined effects of an obesogenic diet and psychological stress on the expression profile and activities of the ROS defense system in the rat brain.

## Methods

Experimental procedures involving rats were performed in compliance with the Loma Linda University School of Medicine regulations and institutional guidelines consistent with the National Institutes of Health Guide for the Care and Use of Laboratory Animals. We made efforts to reduce the number of rats used in the study as well as to minimize animal suffering and discomfort. This study uses *secondary behavioral data analyses* from previously reported findings from our lab as a viable method to clarify the potential involvement of oxidative stress on fear and anxiety-like behavior in rats that were exposed to an obesogenic diet and psychogenic predatory stress [13].

### Animals

Thirty-six (36) adolescent male Lewis rats (postnatal day, PND, 21-23) were acquired from Charles River Laboratories (Portage, MI). The rats were pair-housed and maintained in conventional housing conditions which are, 21 ± 2 °C, relative humidity of 45%, and a 12-hour light/dark cycle with lights turned on at 7:00 am. The animals had *adlibitum* access to food and water.

### Study Design

The rats were allowed to habituate to the animal facility for one week. Afterward, they were divided at random into two groups: one received the control diet (CD; *n* = 18) while the other received the Western-like high-fat diet (WD; *n* = 18). The rats consumed the diets for 8 weeks before exposure to the psychogenic stressor and behavioral testing. During the acclimation/handling sessions, the rats were handled for 5 minutes in their room for 3 consecutive days during the morning hours. Subsequently, the rats were brought to the behavior facility and acclimated to the room for 30 min and then handled for 5 min. Twenty-four hours following the acclimation to the testing environment, the rats were habituated to the Acoustic Startle Reflex (ASR) chamber. This habituation protocol consisted of placing the rats inside the testing enclosure and chamber for 5 min and then gently returning them to their home cage. Following forty-eight hours after habituation, ASR baseline measurements were acquired. Three days after acquiring the baseline ASR measurements, the rats were matched and subdivided into four groups based on exposure to psychogenic stress (control diet unexposed, CDU, *n* = 8; control diet exposed, CDE, *n* = 10; Western high-fat diet unexposed, WDU, *n* = 8; Western high-fat diet exposed, WDE, *n* = 10). We used cat urine as a mild psychogenic stressor. Twenty-four hours after exposure to the mild psychogenic stress, the 6-day fear-potentiated startle (FPS) protocol was commenced. Twenty-four hours after FPS day 6, the rats were tested in the elevated plus-maze (EPM) and ASR. All behaviors were recorded between 10:00-15:00 h. The rats were euthanized 24 h after completion of the behavioral assessments, and plasma and brain tissue were collected.

### Diets

The rats were given *adlibitum* access to water and either standard chow control diet (CD; 13% kcal fats and 53% kcal from carbohydrates; total density = 3.1 kcal; Harlan Laboratories Inc., Indianapolis, IN) or a Western-like high-saturated fat diet (WD; catalog #F6724; 41% kcal fats and 43% kcal from carbohydrates; total density = 4.6 kcal; Bio-Serv, Frenchtown, NJ). The composition of the diets is fully detailed in previous reports (**Table 1)** [13,15].

**Table 1.**
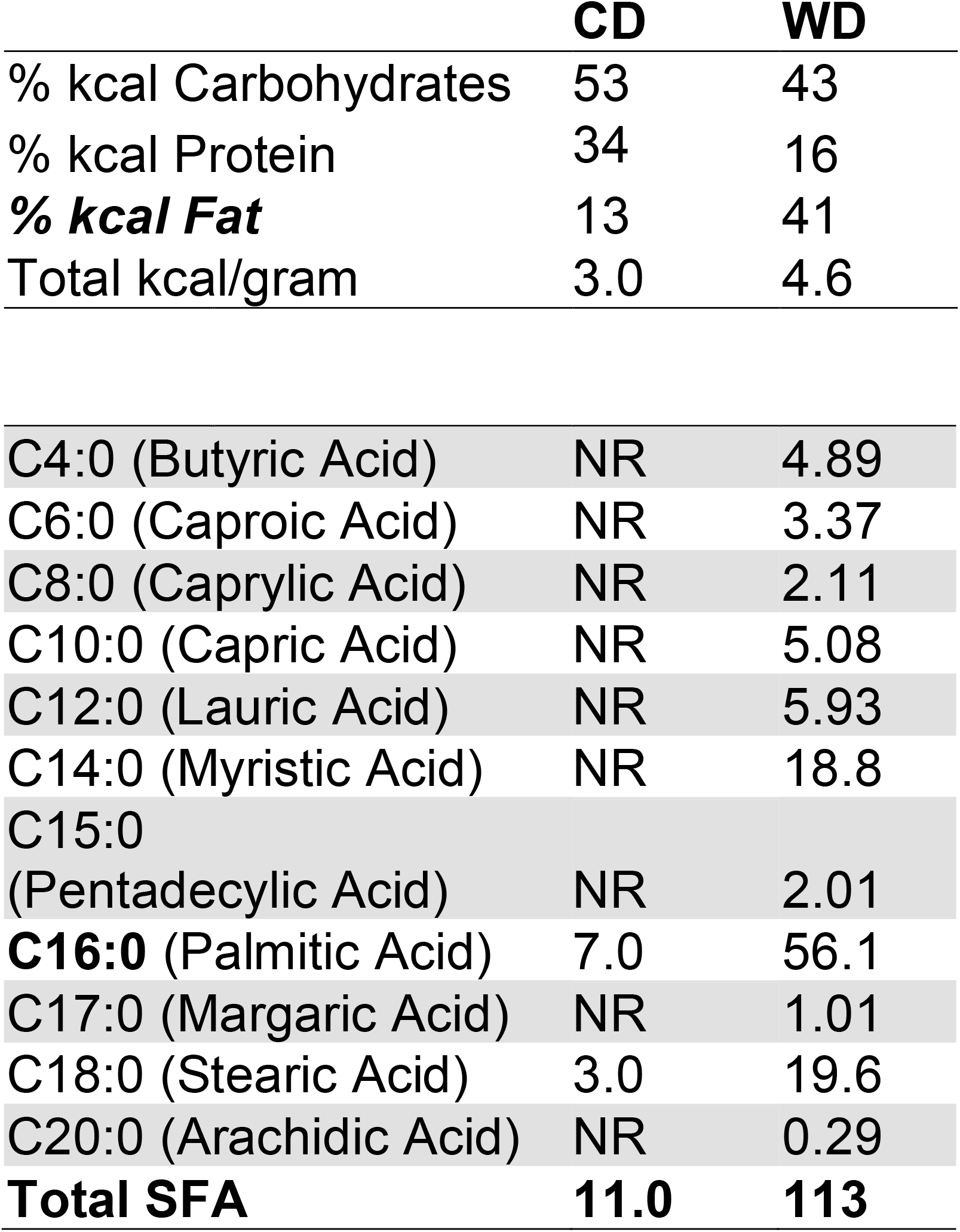
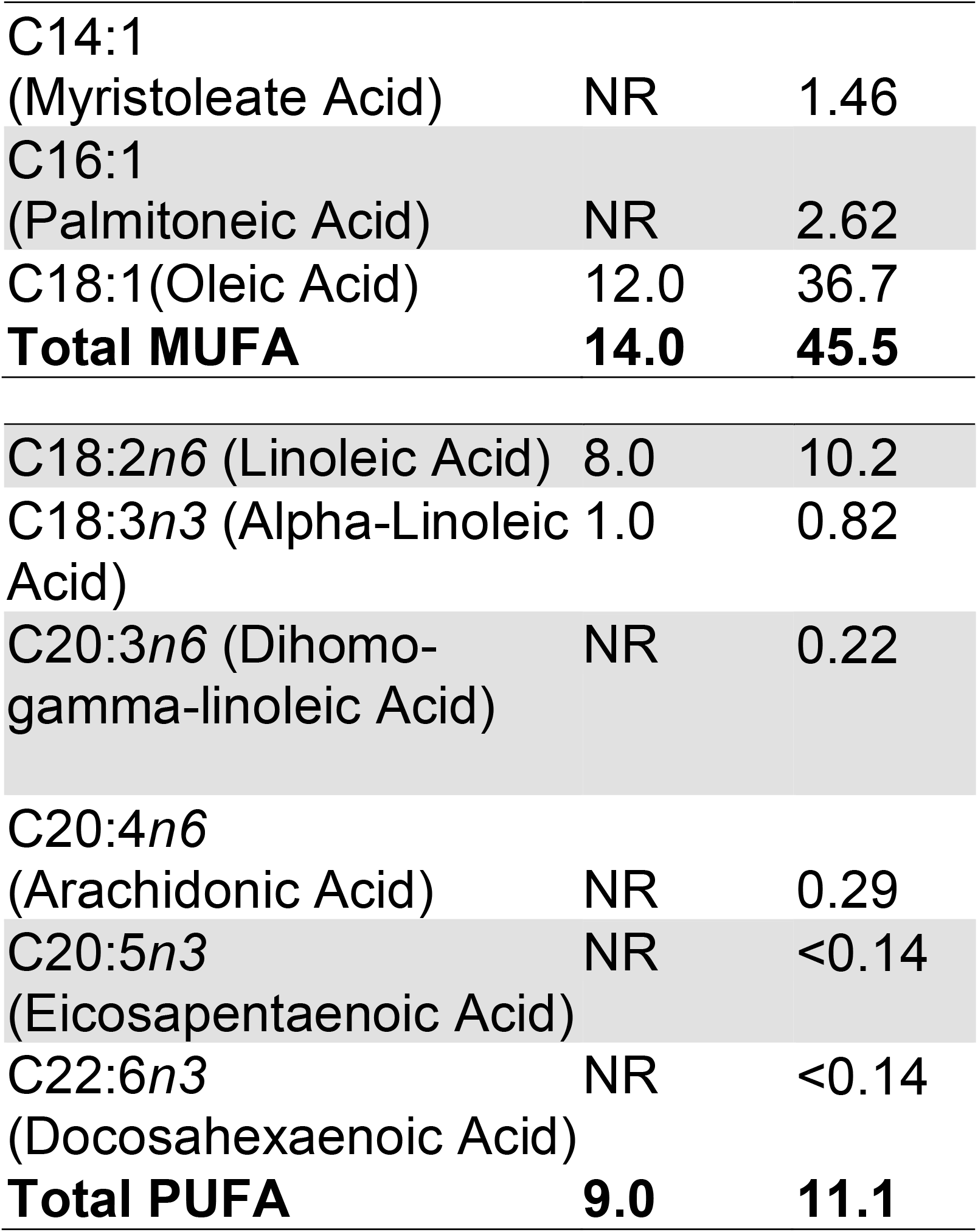
Detailed Diet Composition. CD, control grain-based diet; WD, Western-like high-fat diet

### Predator Odor Stress

Psychogenic stress (PS) was induced by exposing the rats to predator odor [12,13,41,42]. In this model, the rats are exposed to soiled cat litter that was in use by a domestic male cat. The soiled cat litter was placed in small glass containers before the experiments (~100 grams/container). Each rat was placed in an empty standard plastic mouse cage with a filtered plastic top for 15 min. Subsequently, either a clean, fresh litter container (unexposed group) or a soiled cat litter container (exposed group) was placed inside the cage. Each rat was allowed to interact with the container for 15 min. The total time in the cage was 30 min.

### Acoustic startle reflex (ASR)

The ASR experiments were performed using SR-Lab acoustic chambers (San Diego Instruments, San Diego, CA, USA). The ASR magnitudes were measured by placing the rats inside startle enclosures that contained piezoelectric transducers. The acoustic stimuli intensities and response sensitivities were calibrated prior to commencing the experiments. The ASR protocol has been previously described by our group [12,13]. The experimental sessions were 22 min long and started with a 5-min habituation period (background noise = 55 decibels, dB). The rats were then presented with a series of 30 tones (10 tones at each intensity: 90 dB, 95 dB, and 105 dB) using a 30-s inter-trial interval (ITI). The acoustic stimuli had a 20 millisecond (ms) duration, and the trials were presented in a quasi-random order. The rats were returned to their home cages after the completion of the session. The enclosures were cleaned with soap and water and thoroughly dried between sessions. The ASR magnitudes were averaged and normalized to body weight to eliminate confounding factors associated with a differential body weight between groups (Weight-corrected ASR = maximum startle magnitude in mV divided by body weight at testing day) [12,13,43–45].

### Fear potentiated startle (FPS)

The fear-potentiated startle (FPS) protocol was adapted from Dr. Michael Davis [46] and detailed in previous studies from our laboratory [13–15]. Each FPS session started with a 5-min acclimation period (background noise = 55 dB). During the first session of the paradigm (fear training), the rats were conditioned to associate a light stimulus (conditioned stimulus, CS) with a 0.6 mA foot shock (unconditioned stimulus, US). This conditioning session involved 10 CS + US pairing presentations. During each CS + US presentation, the light (3200 ms duration) was paired with a co-terminating foot shock (500 ms duration). Light-shock pairings were presented in a quasi-random manner (ITI = 3-5 min). The acquisition of fear-related responses was measured 24 h later. During this second session (fear acquisition testing), the rats were first presented with 15 startle-inducing tones (*leader trials*; 5 trials for each tone: 90 dB, 95 dB, and 105 dB) delivered alone at 30 sec ITI. Subsequently, the rats were presented with 60 test trials. For half of these test trials, a 20 ms tone was presented alone (**tone alone trials**; 10 trials for each tone: 90 dB, 95 dB, and 105 dB). For the other half, the tone was paired with a 3200 ms light (**light + tone trials**; 10 trials for each pairing: 90 dB, 95 dB, and 105 dB). To conclude the acquisition testing session, the rats were presented with 15 startle-inducing tones (*trailer trials*; 5 trials of each tone: 90 dB, 95 dB, and 105 dB) delivered at 30 sec ITI. The trials in this session were presented in a quasi-random order (ITI = 30 sec). The startle-eliciting tones had a 20 ms duration. One day after fear acquisition testing, the rats were exposed to three (3) consecutive daily fear extinction-training sessions. The fear extinction training session consisted of 30 CS alone presentations (light without shock or noise bursts) with a duration of 3700 ms (ITI = 30 sec). Twenty-four hours after the last extinction session, we determined the FPS responses using a testing session identical to that used to measure the acquisition of fear. The persistence of potentiated startle responses in the presence of the CS during this fear extinction acquisition session indicated a failure to acquire extinction learning. FPS data were reported as the proportional change between US and CS + US [%FPS = ((Light + Tone Startle) − (Tone Alone Startle)) / (Tone Alone Startle) × 100] [47]. Fear extinction was scored as the FPS difference between the fear learning and the fear extinction testing sessions (pre-extinction FPS – post-extinction FPS).

### Elevated Plus Maze (EPM)

The near infrared-backlit (NIR) elevated plus maze (EPM) consisted of two opposite open arms (50.8 × 10.2 × 0.5 cm) and two opposite enclosed arms (50.8 × 10.2 × 40.6 cm) (Med Associates Inc., Fairfax, VT). The arms were connected by a central 10 × 10 cm square-shaped area. The maze was elevated 72.4 cm from the floor. Behaviors were recorded in the dark. The rats were placed on the central platform facing an open arm and allowed to explore the EPM for 5 min. The apparatus was cleaned after each trial (70% ethanol, rinsed with water, and then dried with paper towels). The behaviors were monitored via a monochrome video camera equipped with a NIR filter and recorded and tracked using Ethovision XT (Noldus Information Technology, Leesburg, VA). In rats, changes in the percentage of time spent on the open arms (OA) indicate changes in anxiety, and the number of closed arms (CA) entries is the best measure of locomotor activity [48,49]. These data are used to calculate the anxiety index [12,50–52].

Anxiety Index = 1 − [([OA cumulative duration / Total test duration] + [OA entries / Total number of entries to CA + OA]) / 2].

### Immunoblotting

A group of rats (*n* = 24; 6 rats per group) were deeply anesthetized with an intraperitoneal overdose injection of Euthasol (150 mg/kg; Virbac, Fort Worth, TX) and perfused transcardially with PBS. Following perfusions, the rats were rapidly decapitated, and the brains isolated. The brain tissue was homogenized and transferred to a 1.5 mL microtube containing 700 μl of cold Cell Lytic MT Lysis Extraction Buffer (catalog #C3228; Sigma-Aldrich, St. Louis, MO), 1% phosphatase inhibitor cocktail 3 (catalog #P0044; Sigma-Aldrich), and a FAST Protease Inhibitor Cocktail Tablet, EDTA free (catalog #S8830; Sigma-Aldrich). Protein extraction was performed as suggested by the manufacturer. Briefly, the samples were centrifuged for 10 min at 4°C (20,000 rpm), and the supernatant was collected and stored at −80°C until needed for further processing. As for protein quantification, the Bio-Rad protein assay was used according to the manufacturer’s instructions (Bio-Rad Laboratories, Hercules, CA). Proteins were separated on a 12% polyacrylamide-SDS gel (40 μg of protein/lane) and wet transferred to a nitrocellulose membrane from 1h at 4°C. The membrane was blocked with Odyssey Blocking Buffer (catalog #927-40000; LI-COR Biosciences, Lincoln, NE, USA) for 1 hour at room temperature. The proteins of interests were detected using the following primary antibodies: anti-rabbit Catalase (1:375; catalog #Ab52477; Abcam, Cambridge, MA); anti-rabbit Glutathione Peroxidase (1:500; catalog #Ab22604; Abcam); anti-rabbit Glutathione Reductase (1:2000; catalog #Ab16801; Abcam); anti-rabbit SOD-1 (1:500; catalog #Ab16831; Abcam); anti-rabbit SOD-2 (1:2000; catalog #13141S; Cell Signaling, Danvers, MA); anti-rabbit GFAP (1:5000; cat# 556,327, BD Biosciences, San Jose, CA, USA); and anti-rabbit Iba-1 (1:1000; cat# 016-20,001 Wako Chemicals, Richmond, VA, USA). All the antibodies were incubated in a blocking solution and overnight at 4°C. Anti-mouse ß-actin (1:10000; catalog #A5441; Sigma-Aldrich, St. Louis, MO) was used as a loading control for antioxidant proteins. GFAP and Iba-1 protein analyses were generated from previously collected data and normalized to GAPDH (1:5000; cat# G9545, Sigma-Aldrich) [13]. As for secondary antibodies, we used goat anti-rabbit (1:25000; catalog #926-32211; LI-COR Biosciences) and goat anti-mouse (1:25000; catalog #925-68070; LI-COR Biosciences) for 1h at room temperature. Infrared signals from membranes were detected using the Odyssey CLx Scanner (LI-COR Biosciences, Lincoln, NE). Densitometric analyses were performed using the Image Studio 5.2 Software (LI-COR Biosciences, Lincoln, NE).

### Glutathione Peroxidase and Reductase Activity

The plasma glutathione peroxidase (GPx) and reductase (GSHR) activities were determined using a Glutathione Peroxidase Assay Kit (#703102) and a Glutathione Reductase Assay Kit (#703202; Cayman Chemical Company, Ann Arbor, MI) according to the manufacturer’s instructions. The rate of change of absorbance (optical density at 340 nm) per minute was measured using the SpectraMax® i3x Multi-Mode Microplate Reader (Molecular Devices, Molecular Devices, Sunnyvale, CA, USA) and expressed as nmol/min/ml.

### Statistical Analysis

We recorded and tabulated all data using Excel (Microsoft, Redmond, WA, USA). Statistical analyses were performed using Graphpad Prism version 8.0 (Graphpad Software, La Jolla, CA, USA). Main (diet, stress) and interaction (diet × stress) effects were determined with two-way analysis of variance (ANOVA) followed by Tukey’s post hoc test. To explore associations between behavioral phenotypes and ROS biomarkers, we used Pearson’s correlation test. We considered differences significant if *p* < 0.05; all data are shown as the mean ± SEM.

## Results

This study investigated the relative and combined effects of the obesogenic Western-like diet (**WD**) and psychogenic stress (**PS**) on anxiety-like behaviors and neuroinflammation/oxidative stress biomarkers. **Figure 1** illustrates the experimental timeline and procedures (adapted from Vega-Torres et al., 2018) [13]. Body weight, food consumption, and obesogenic biomarker data was previously reported [13]. We found that the exposure to the psychogenic stressor increased the reactivity to the footshocks during the fear conditioning session (FPS day 1) [*F*_(1,32)_ = 6.27, *p* = 0.02] (**Figure 2A**). Analyses revealed no significant effects of the diet type [*F*_(1,32)_ = 0.06, *p* = 0.81] or interaction between factors [*F*_(1,32)_ = 1.02, *p* = 0.32] on footshock reactivity. We investigated the effect of the obesogenic diet and psychogenic stress in fear-related behaviors. As previously reported, analysis of the effect of diet revealed that the obesogenic WD significantly reduced fear learning, as evaluated by the fear-potentiated startle [*F*_(1,31)_ = 10.80, *p* = 0.003] (**Figure 2B**). Psychogenic stress exposure [*F*_(1,32)_ = 0.56, *p* = 0.46] and interaction between the obesogenic diet and PS [*F*_(1,32)_ = 0.15, *p* = 0.70] did not have a significant effect on fear acquisition. Analysis of the effect of the diet on fear extinction learning revealed a significantly attenuated fear extinction index in the rats that consumed the obesogenic diet [*F*_(1,32)_ = 10.57, *p* = 0.003] (**Figure 2C**). The psychogenic stressor [*F*_(1,32)_ = 0.01, *p* = 0.91] and interaction [*F*_(1,32)_ = 0.04, *p* = 0.81] did not affect fear extinction significantly.

**Figure 1.**
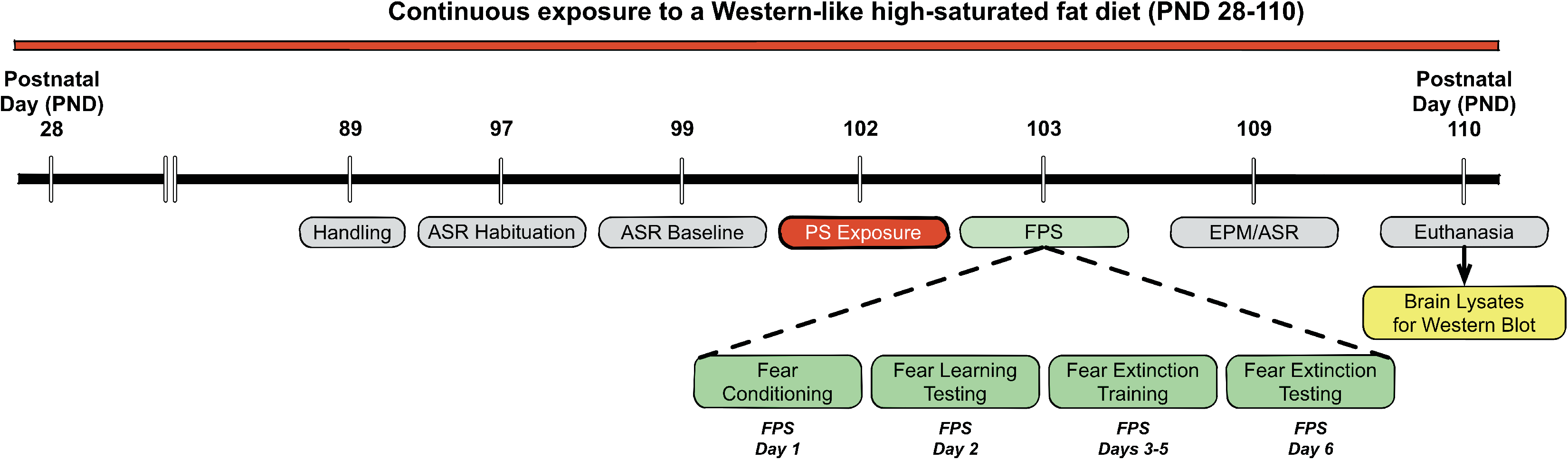
Study design and timeline of experimental procedures, behavioral tests, and outcome measures. Abbreviations: PND, postnatal day; CDU, control diet unexposed; CDE, control diet exposed; WDU, Western high-fat diet unexposed; WDE, Western high-fat diet exposed; ASR, acoustic startle reflex; EPM, elevated plus maze; FPS, Fear Potentiated Startle.

**Figure 2.**
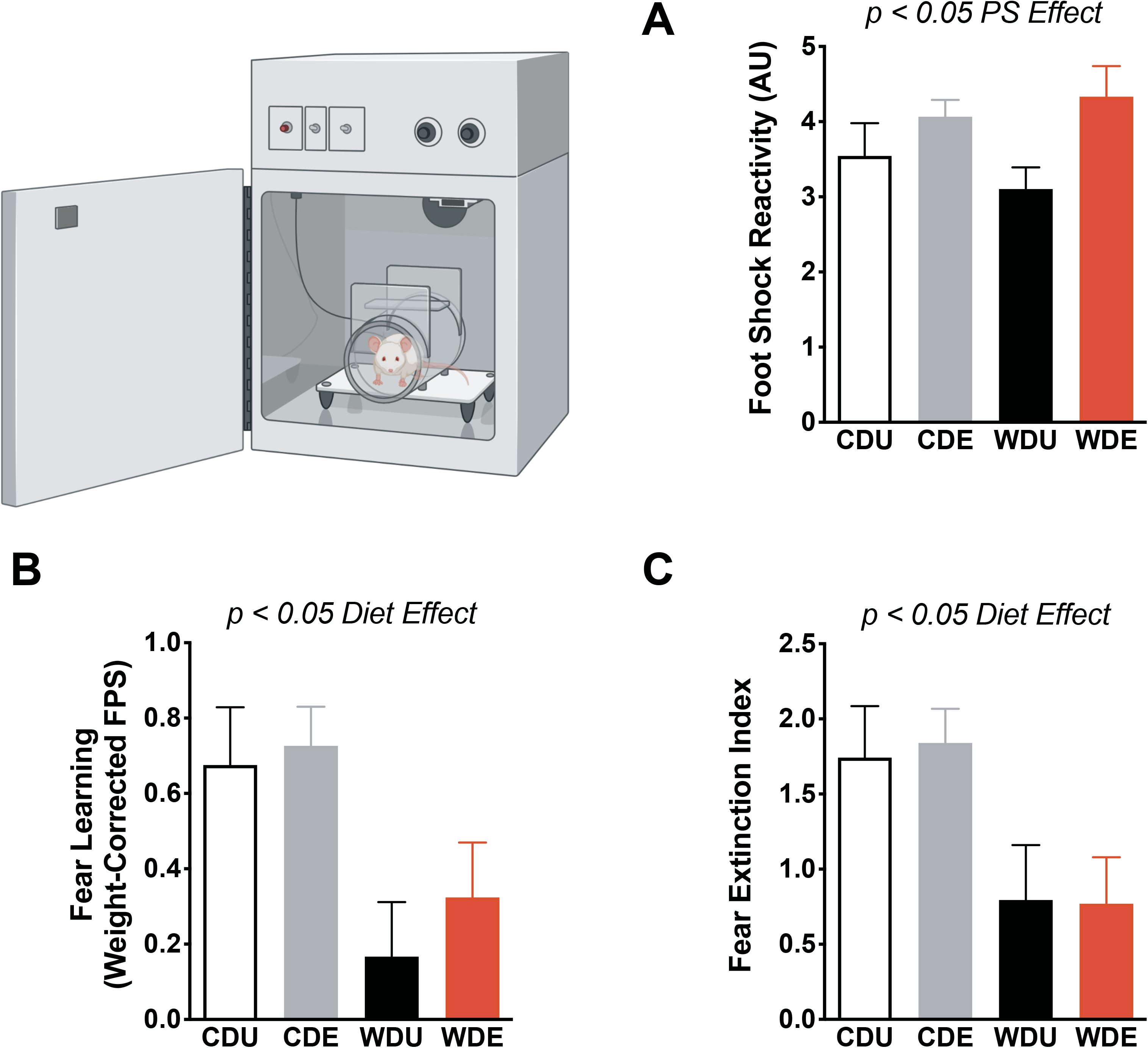
Deficits in delay cued fear conditioning and fear extinction in rats that consume an obesogenic Western-like diet. **(A)** Psychogenic stress (PS) significantly increase footshock reactivity during the fear-conditioning session [stress: *F*_(1,32)_ = 6.27, *p* = 0.018]. **(B)** Average fear-potentiated startle (FPS) responses 24 h after conditioning. The rats that consumed the obesogenic Western-like diet (WD) exhibited reduced fear learning relative to controls [diet: *F*_(1,31)_ = 10.80, *p* = 0.003]. **(C)** Average extinction index following three consecutive daily fear extinction-training sessions. WD rats displayed deficits in fear extinction [diet: *F*_(1,32)_ = 10.57, *p* = 0.003]. *, *p* < .05; *n* = 8-10 rats/group. Error bars are S.E.M.

Next, we tested whether the obesogenic WD, PS, or interactions between these factors had a significant effect on anxiety-like behaviors in the elevated plus maze (EPM). Exposure to the PS increased general measures of anxiety in the EPM [*F*_(1,32)_ = 5.17, *p* = 0.03] (**Figure 3A**). The diet type [*F*_(1,32)_ = 0.08, *p* = 0.78] and interactions between factors [*F*_(1,32)_ = 0.78, *p* = 0.38] did not have significant effects on the anxiety index. We found no significant main effects or interactions on the number of open arm entries (diet: *F*_(1,32)_ = 0.89, *p* = 0.77; PS: *F*_(1,32)_ = 2.91, *p* = 0.10; interaction: *F*_(1,32)_ = 32.33, *p* = 0.57) (**Figure 3B**). Analyses revealed that rats exposed to the PS exhibited reduced duration in the open arms of the EPM [*F*_(1,32)_ = 6.73, *p* = 0.01] (**Figure 3C**). We found no significant diet [*F*_(1,32)_ = 0.14, *p* = 0.71] and interaction [*F*_(1,32)_ = 1.23, *p* = 0.28] effects on the duration spent in the open arms. Analyses showed no significant main effects or interactions on the total distance traveled in the EPM (diet: *F*_(1,32)_ = 1.18, *p* = 0.29; PS: *F*_(1,32)_ = 0.82, *p* = 0.37; interaction: *F*_(1,32)_ = 0.12, *p* = 0.73) (**Figure 3D**). PS exposure reduced nose duration in head-dipping zones of the EPM [*F*_(1,32)_ = 5.43, *p* = 0.03], supporting the anxiogenic effect of the stressor (**Figure 3E**). Post hoc analyses revealed that the CD rats exposed to PS exhibited a significantly reduced duration in head-dipping zones relative to unexposed CD rats (*p* = 0.04). The diet type [*F*_(1,32)_ = 1.68, *p* = 0.20] and interactions [*F*_(1,32)_ = 2.51, *p* = 0.12] did not have a significant effect on the duration of the rats in head-dipping zones. PS increased the frequency of ethological measures of anxiety in the EPM. We found that both the frequency of stretch attend postures (PS: *F*_(1,32)_ = 5.26, *p* = 0.03; diet: *F*_(1,32)_ = 0.66, *p* = 0.42; interaction: *F*_(1,32)_ = 1.29, *p* = 0.26) (**Figure 3F**) and unsupported rearing (PS: *F*_(1,31)_ = 7.42, *p* = 0.01; diet: *F*_(1,31)_ = 1.06, *p* = 0.31; interaction: *F*_(1,31)_ = 0.28, *p* = 0.60) (**Figure 3G**) were significantly increased in PS-exposed rats. The time spent performing spontaneous self-grooming behaviors in the EPM was not significantly affected by the diet, PS, or the interaction between these factors (diet: *F*_(1,31)_ = 0.003, *p* = 0.96; PS: *F*_(1,31)_ = 0.60, *p* = 0.44; interaction: *F*_(1,31)_ = 2.09, *p* = 0.16) (**Figure 3H**).

**Figure 3.**
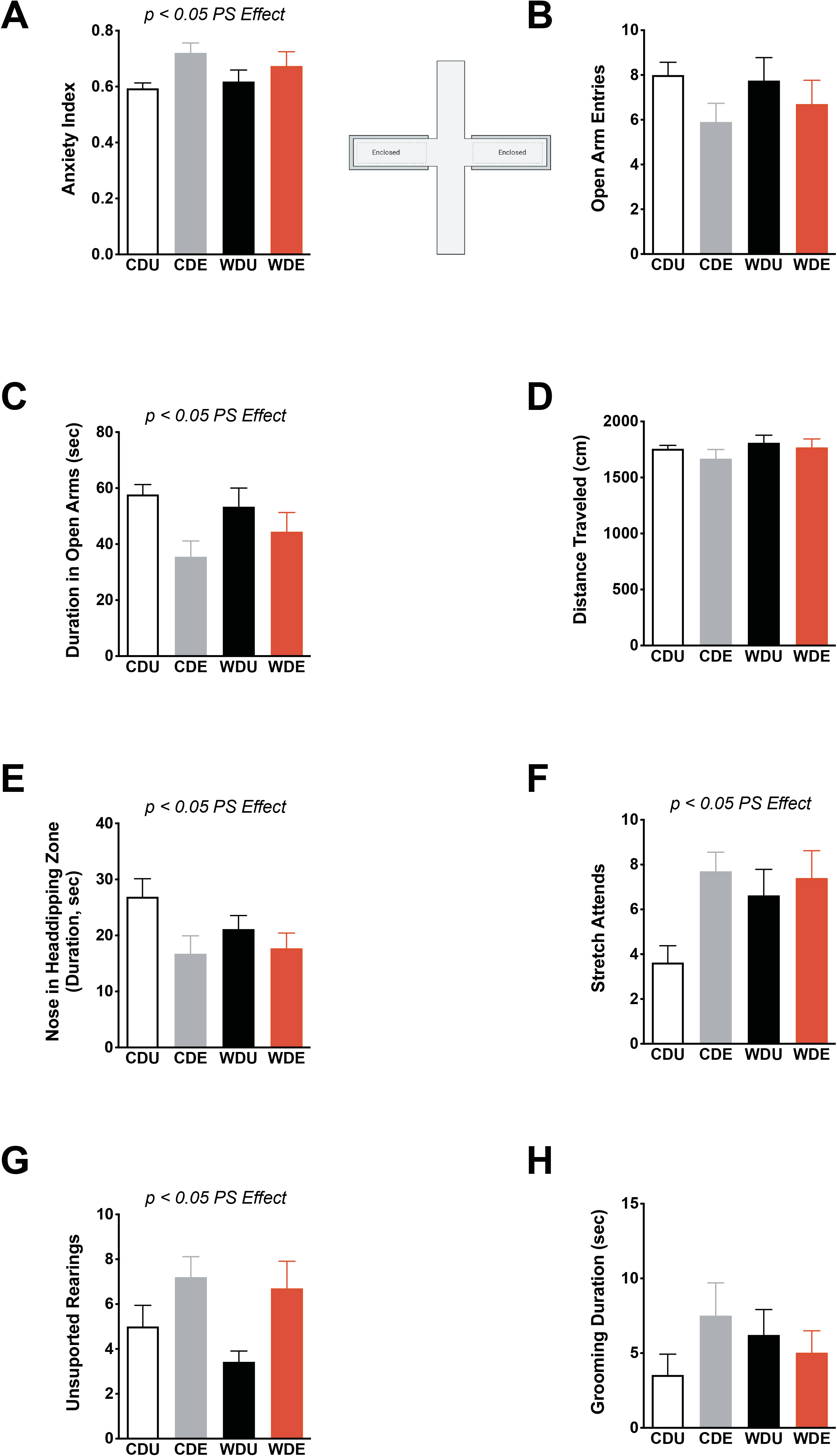
Selective anxiogenic effects of predatory odor stress in the elevated plus maze. **(A)** Average anxiety index scores calculated from behavioral outcomes in the elevated plus maze (EPM). PS heightened anxiety-like behaviors in the EPM [stress: *F*_(1,32)_ = 5.17, *p* = 0.030]. **(B)** Average open arm entries were similar between groups (*p* > 0.05). **(C)** Average duration in the open arms. PS rats exhibited reduced duration in the open arms of the EPM [stress: *F*_(1,32)_ = 6.73, *p* = 0.014]. **(D)** Average distance traveled was similar between groups (*p* > 0.05). **(E)** Average duration of nose in head-dipping zones of the EPM, expressed in seconds. PS significantly decreased the extent of nose in head-dipping zones [stress: *F*_(1,32)_ = 5.43, *p* = 0.026]. **(F)** Average frequency of stretch attend postures (SAP). PS increased SAP frequency [stress: *F*_(1,32)_ = 5.26, *p* = 0.030]. **(G)** Average frequency of unsupported rearing behaviors during EPM testing. PS increased the frequency of unsupported rearings [*F*_(1,31)_ = 7.42, *p* = 0.011]. **(H)** Average grooming duration (in seconds) was similar between groups (*p* > 0.05). *n* = 8-10 rats/group. Error bars are S.E.M.

To examine the adaptive responses of the brain antioxidant network to the obesogenic WD and PS, we focused on measuring the expression levels of critical proteins implicated with oxidative stress (**Figure 4A**). We found that the WD had a robust effect on the expression of catalase in the brain (diet: *F*_(1,18)_ = 8.60, *p* = 0.01; PS: *F*_(1,18)_ = 2.84, *p* = 0.11; interaction: *F*_(1,18)_ = 0.47 *p* = 0.50) (**Figure 4B**). Post hoc revealed significantly increased catalase protein expression levels in the brain of WDE rats relative to CDU (*p* = 0.03), indicating that the obesogenic WD exacerbates the effect of PS on catalase protein expression. We found no significant main or interaction effects on the protein expression levels of glutathione peroxidase (diet: *F*_(1,18)_ = 3.08, *p* = 0.10; PS: *F*_(1,18)_ = 0.002, *p* = 0.10; interaction: *F*_(1,18)_ = 0.050, *p* = 0.83) (**Figure 4C**), glutathione reductase (diet: *F*_(1,17)_ = 0.32, *p* = 0.58], PS: *F*_(1,17)_ = 0.81, *p* = 0.38; interaction: *F*_(1,17)_ = 0.31, *p* = 0.58) (**Figure 4D**), superoxide dismutase-1 (diet: *F*_(1,18)_ = 0.01, *p* = 0.91; PS: *F*_(1,18)_ = 0.03, *p* = 0.87; interaction: *F*_(1,18)_ = 0.02, *p* = 0.88) (**Figure 4E**), and superoxide dismutase-2 (diet: *F*_(1,19)_ = 0.53, *p* = 0.48; PS: *F*_(1,19)_ = 0.20, *p* = 0.66; interaction: *F*_(1,19)_ = 0.81, *p* = 0.38) (**Figure 4F**).

**Figure 4.**
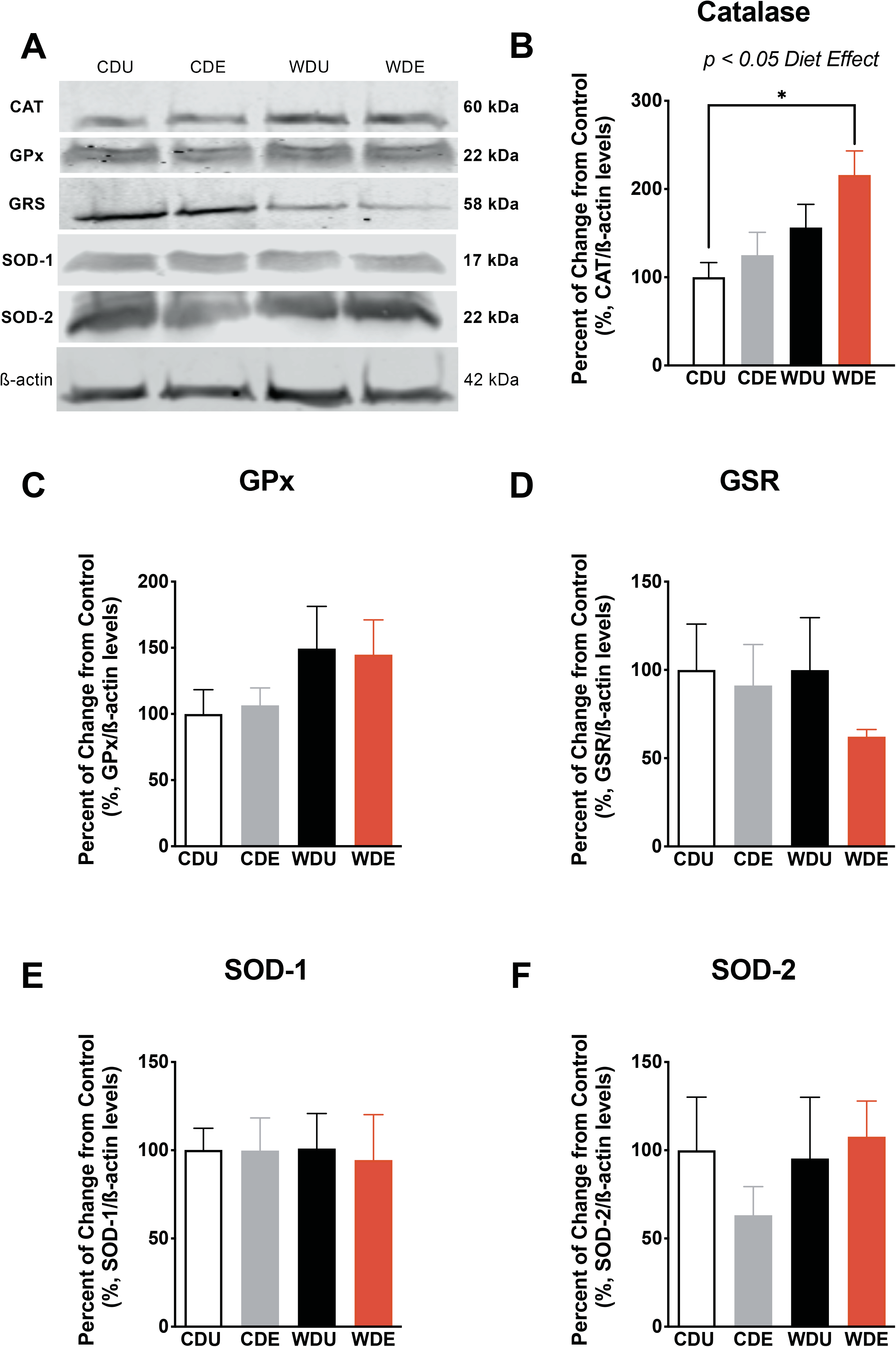
Increased catalase protein levels in the brain of rats exposed to an obesogenic diet. **(A)** Representative Western blot bands from whole-brain homogenates illustrating the relative levels of critical proteins involved in the brain antioxidant network. Protein levels are expressed in percent from control and normalized in relation to **β**-actin. **(B)** The obesogenic WD increased catalase protein levels in the brain [diet: *F*_(1,18)_ = 8.60, *p* = 0.0090]. This effect was particularly evident in WDE rats relative to CDU (post-hoc *p* = 0.03), indicating that PS exacerbates the effect of an obesogenic diet on catalase protein levels. The protein levels of glutathione peroxidase **(C),** glutathione reductase **(D)**, superoxide dismutase 1 **(E)**, and superoxide dismutase 2 **(F)** were not significantly affected by the dietary and stress manipulations (for main and interaction effects: *p* > 0.05). *n* = 5-6 rats/group. Error bars are S.E.M.

Astrocytes and microglia are resident immune-competent cells with high expression of catalase and important roles regulating oxidative stress [53–56]. With the ultimate aim of identifying whether the obesogenic WD and PS impacted these relevant cellular phenotypes, we next determined the expression levels of GFAP and Iba-1 via Western blot (**Figure 5A)**. In agreement with previous reports from our group, we found that the obesogenic diet increased GFAP protein levels in the brain (diet: *F*_(1,13)_ = 16.50, *p* = 0.001; PS: *F*_(1,13)_ = 0.59, *p* = 0.46; interaction: *F*_(1,13)_ = 2.54, *p* = 0.14) (**Figure 5B**). Post hoc analyses demonstrated that WDE rats had increased GFAP levels relative to CDU (*p* = 0.02) and CDE (*p* < 0.01) rats. Interestingly, we found no significant main or interaction effects on the expression of Iba-1 in the brain (diet: *F*_(1,15)_ = 0.045, *p* = 0.83; PS: *F*_(1,15)_ = 0.25, *p* = 0.63; interaction: *F*_(1,15)_ = 0.24, *p* = 0.63 (**Figure 5C**), revealing a selective effect of the experimental manipulations on glial markers. We found strong associations between the glial markers and catalase levels in the brain. Notably, while GFAP levels were negatively associated with catalase protein expression levels (Pearson’s *r* = −0.68, *p* = 0.03) (**Figure 5D**), Iba-1 protein levels were positively associated (Pearson’s *r* = 0.89, *p* = 0.004) (**Figure 5E**).

**Figure 5.**
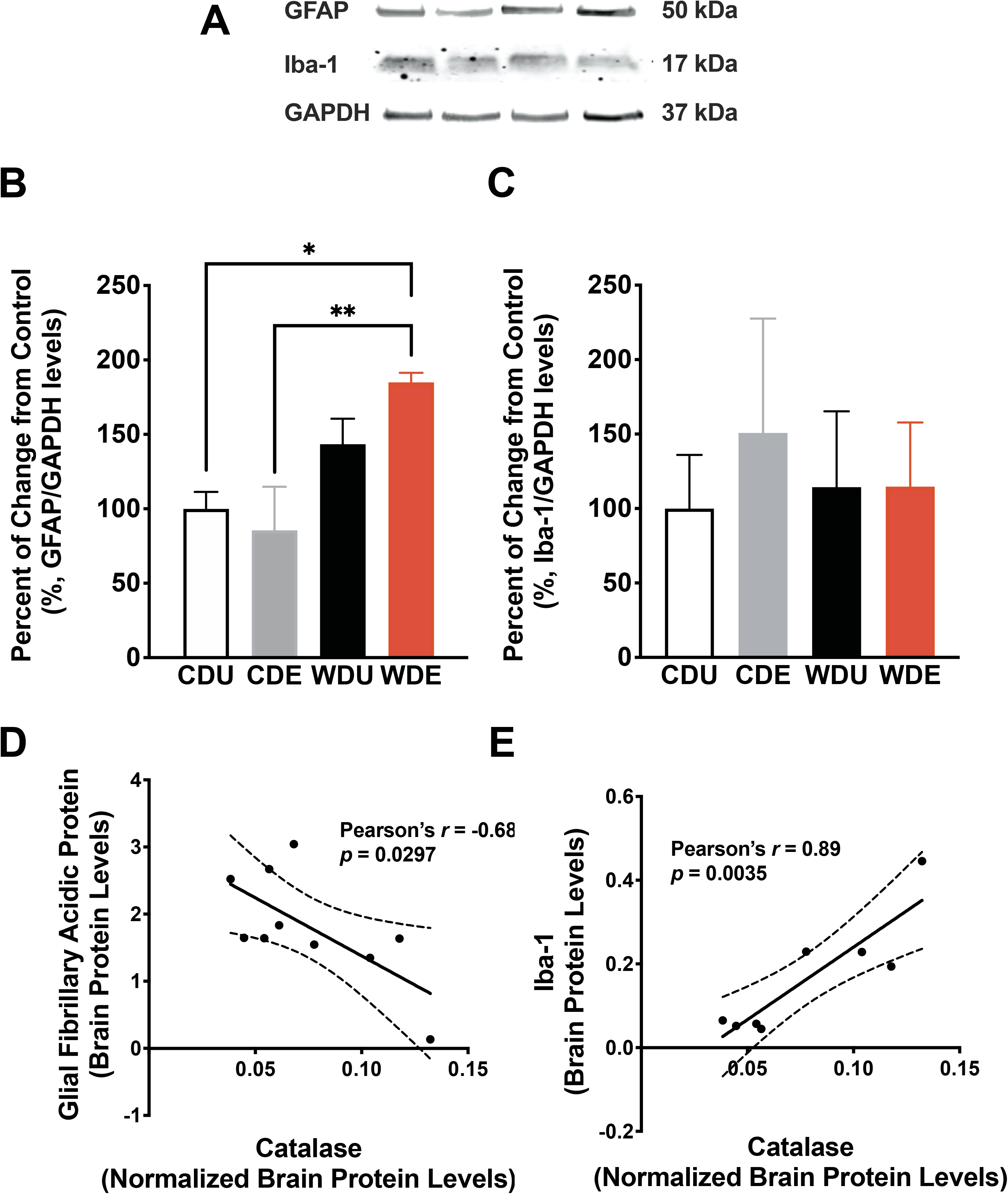
Psychogenic stress synergizes with an obesogenic WD to increase GFAP protein levels. **(A)** Western blot data from Vega-Torres et al. 2018 was reanalyzed to include the effects of PS on neuroinflammation [13]. **(B)** The obesogenic WD increased GFAP (glial fibrillary acidic protein) levels in whole-brain homogenates [*F*_(1,13)_ = 16.50, p = 0.0013]. Interestingly, analyses demonstrate that WDE rats had increased GFAP levels relative to CDU (*p* = 0.02) and CDE (*p* < 0.01) rats. **(C)** WD and PS exposure did not alter Iba-1 (Ionized calcium-binding adaptor molecule 1) protein levels (*p* > 0.05). *n* = 5-6 rats/group. Error bars are S.E.M. Protein levels are expressed in percent from control and normalized to GAPDH. **(D)** A scatter plot shows a significant inverse relationship between GFAP and catalase protein levels in the brain (Pearson’s *r* = −0.68; *p* = 0.030). **(E)** A scatter plot illustrates a robust positive association between Iba-1 and catalase levels in the brain (Pearson’s *r* = 0.89; *p* = 0.0035).

We then determined the enzymatic activity of glutathione peroxidase and reductase in plasma to more clearly define the contributions of the obesogenic WD and PS on peripheral biomarkers of oxidative stress status. Our analyses demonstrate that the obesogenic WD increased the activities of glutathione reductase in plasma (diet: *F*_(1,19)_ = 5.08, *p* = 0.04; PS: *F*_(1,19)_ = 0.79, *p* = 0.38; interaction: *F*_(1,19)_ = 1.12, *p* = 0.30) (**Figure 6A**). On the other hand, we found no significant main or interaction effects for the enzymatic activity of glutathione peroxidase in the plasma (diet: *F*_(1,19)_ = 0.02, *p* = 0.90; PS: *F*_(1,19)_ = 1.17, *p* = 0.29; interaction: *F*_(1,19)_ = 1.09, *p* = 0.77). Given that peripheral ROS can influence brain oxidative status , we investigated associations between the enzymatic activities of glutathione peroxidase and reductase and catalase protein levels in the brain. Pearson’s correlation analyses revealed a robust negative association between glutathione reductase activity in plasma and catalase protein expression in the brain (Pearson’s *r* = −0.79, *p* = 0.002) (**Figure 6C**). We also found a significant relationship between glutathione peroxidase activity in plasma and catalase protein expression in the brain (Pearson’s *r* = −0.68, *p* = 0.04) (**Figure 6D**).

**Figure 6.**
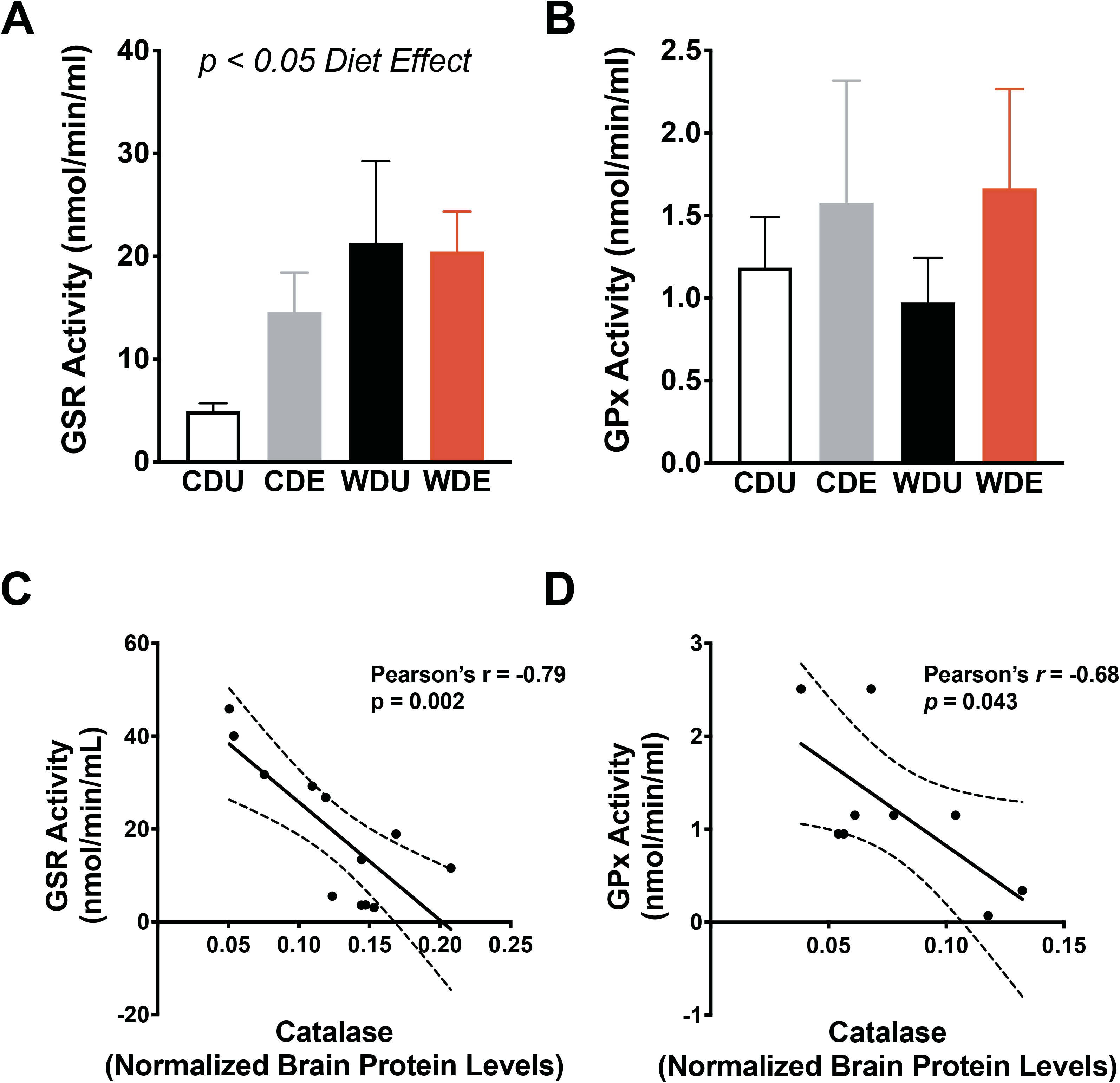
Increased plasma glutathione reductase activities in rats exposed to an obesogenic diet. **(A)** The obesogenic WD increased the averaged glutathione reductase enzymatic activity in plasma [*F*_(1,19)_ = 5.08, *p* = 0.036]. **(B)** Average plasma glutathione peroxidase activities were similar between diet groups (*p* > 0.05). For A and B: *n* = 5-6 rats/group, error bars are S.E.M. **(C)** Scatter plot shows a significant inverse relationship between GSR plasma activity and catalase protein levels in the brain (Pearson’s *r* = −0.79; *p* = 0.0020). **(D)** A scatter plot illustrates a significant inverse association between GPx plasm activity and catalase levels in the brain (Pearson’s *r* = 0.68; *p* = 0.0043).

Finally, to investigate correlations between behavioral outcomes and neuroinflammation/oxidative stress biomarkers, we performed Pearson’s correlation analyses. Results revealed significant and relevant associations between behavioral outcomes implicated with fear and anxiety and inflammation/ROS biomarkers not only in CD rats (**Table 2**) but also when analyzing all the rats used in this study (**Table 3**).

**Table 2.** Associations between neuroinflammatory and oxidative stress biomarkers and behavioral outcomes in rats fed the control diet (CDU and CDE groups included).

| NEUROINFLAMMATION/<br>OXIDATIVE STRESS<br>MARKER | BEHAVIOR/<br>BIOCHEMICAL<br>MARKER | P-VALUE | PEARSON'S R |
| --- | --- | --- | --- |
| Catalase | GFAP | 0.030 | -0.682 |
| Catalase | GPx Activity | 0.043 | -0.682 |
| Catalase | Iba-1 | 0.003 | 0.885 |
| Catalase | SOD-2 | 0.009 | 0.807 |
| GFAP | Catalase | 0.030 | -0.682 |
| GFAP | GPx Activity | 0.034 | 0.705 |
| GFAP | Iba-1 | 0.005 | -0.871 |
| Iba-1 | Catalase | 0.003 | 0.885 |
| Iba-1 | GFAP | 0.005 | -0.871 |
| Iba-1 | Open Arm Duration | 0.045 | 0.677 |
| SOD-2 | Catalase | 0.009 | 0.807 |
| SOD-2 | Fear Learning | 0.025 | 0.732 |
| SOD-2 | Extinction Index | 0.018 | -0.658 |
| SOD-2 | Anxiety Index | 0.025 | 0.731 |
| SOD-2 | Stretch Attends | 0.008 | 0.814 |
| SOD-2 | Head Dips | 0.017 | 0.762 |
| SOD-2 | Open Arm Duration | 0.007 | 0.817 |
| SOD-2 | Corticosterone | 0.038 | -0.781 |
| SOD-2 | Body Weight | 0.015 | 0.771 |
| SOD-2 | Footshock Reactivity | 0.017 | -0.760 |

**Table 3.**
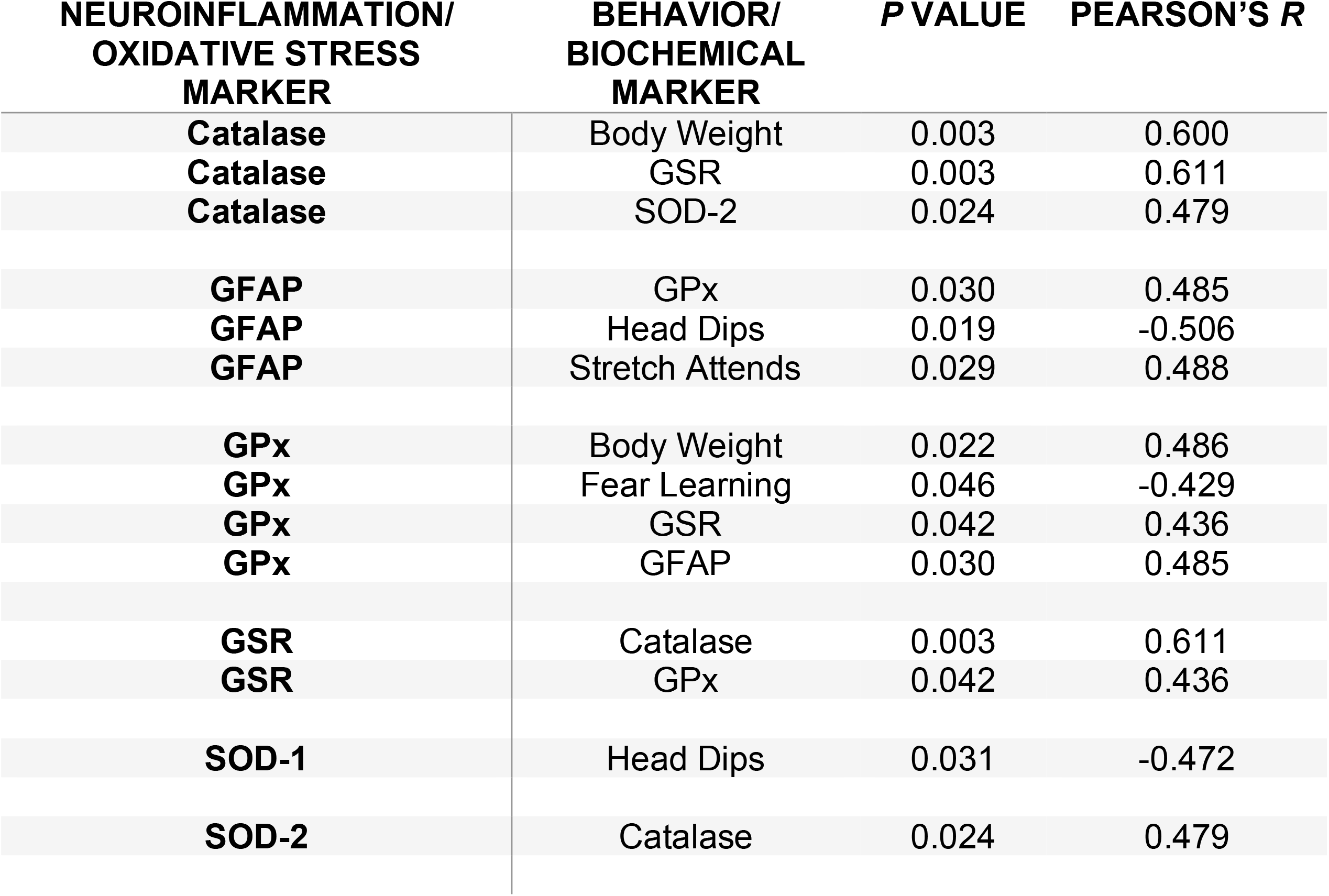
Associations between neuroinflammatory and oxidative stress biomarkers and behavioral outcomes in all study groups.

## Discussion

This study investigated the individual and combined effects of a Western-like high-saturated fat obesogenic diet (WD) and acute exposure to predator odor stress (PS) on critical endogenous antioxidants and redox regulators. The main finding of this study is that WD intake and PS exposure acted synergistically to increase the brain protein levels of specific redox/neuroinflammatory biomarkers. In particular, the protein levels of catalase (CAT) and the glial fibrillary acidic protein (GFAP). We demonstrate that the levels and activities of some of these biomarkers are closely associated with behavioral proxies related to fear and anxiety in rats.

### Obesogenic diets trigger redox dysregulation and neuroinflammation: Implications for stress-related disorders

Obesity is highly comorbid with anxiety and stress-related disorders. Our studies indicate that consumption of an obesogenic WD during adolescence heightens stress reactivity, promotes anxiety-like behaviors, and impairs the structural integrity of neural substrates implicated in stress-induced psychopathology in rats [12–15]. Furthermore, there is support to the notion that impairments in brain and behavior can occur even in the absence of an obesogenic phenotype [13,57–63]. Yet, the molecular players underlying these abnormalities remain unclear. There is mounting evidence demonstrating that diet-induced obesity (DIO) triggers oxidative brain damage and mitochondrial dysfunction in critical corticolimbic regions implicated with stress-related disorders [27,64,65].

Paralleling studies in obese humans [66,67], a large body of evidence indicates that dietary obesity leads to increased production of ROS while altering the brain antioxidant capacity in rats [65]. Notably, DIO-resistant rats exhibit normal ROS levels and no oxidative brain damage, even after the consumption of a high-fat obesogenic diet [27]. This finding suggests that the genetic background strongly influences dietary obesity and oxidative brain damage.

Emerging evidence suggests that the molecular consequences of PTSD include elevated systemic levels of oxidative stress and inflammation [68,69]. Clinical studies have reported significant differences in blood antioxidant enzyme concentrations and ROS-related gene expression between PTSD patients and controls [68,70]. Animal studies have also found evidence consistent with an association between stress reactivity and ROS markers [69]. For example, studies demonstrate the potential involvement of ROS in rat models of PTSD [69,71]. Brain nitric oxide plays a critical role in facilitating fear conditioning [72]. In support of these observations, a study published by Mejia-Carmona et al. showed that predatory stress induces changes in oxidative stress biomarkers in the amygdala and prefrontal cortex in rats [73,74]. Furthermore, ROS heightens stress responsivity and alters fear and anxiety responses in rats [32,75–77]. Interestingly, the PTSD-like rats that were treated with a vitamin E-based antioxidant therapy exhibited resilience to the effects of stress on cognitive function [71]. While we did not detect changes in redox status biomarkers following PS exposure, we identified significant associations between several behaviors implicated in fear and stress reactivity and the protein levels of several ROS status biomarkers. The discrepancies between studies may be related to the mild psychogenic stressor used in this study, a potential masking effect of footshock stress, and the rat strain used in this study. For instance, Lewis rats display higher ROS levels relative to the Brown Norway rat strain, which is protective against Toxoplasma infection resistance [78]. Notably, Lewis rats are highly susceptible to inflammatory challenges and stress [79–81]. Overall, these studies indicate that obesity and the consumption of obesogenic diets rich in saturated fats could affect the mitochondrial redox balance and contribute to brain structural abnormalities and stress-related psychopathology. It is plausible that the consumption of obesogenic diets may “prime” the brain for heightened stress reactivity through increased ROS levels. Thus, ROS levels may represent a risk factor for anxiety and stress-related disorders.

An interesting finding of this study is the identification of selective neuroinflammatory and oxidative stress-related responses to an obesogenic WD and PS. In particular, catalase (CAT) protein levels were upregulated in the brain of the rats that consumed the obesogenic diet and were exposed to the psychogenic stressor. CAT is one of the major endogenous antioxidant enzymes that detoxify the reactive oxygen species (ROS) hydrogen peroxide (H_2_O_2_) to water and oxygen. While brain CAT activity is low compared to other tissues and organs [25], this enzyme regulates macrophage polarization in adipose tissue [82]. Overexpression of catalase is beneficial in the context of obesity. For instance, overexpression of a mitochondria-specific catalase prevents insulin impairments and inflammation while attenuating mitochondrial ROS production in mice on a brief high-fat diet [83].

On the other hand, catalase deletion leads to metabolic impairments and an obesogenic phenotype in mice [84]. Interestingly, it has been demonstrated that the reaction of CAT to dietary fat can occur before the manifestation of obesity-related phenotypes in mice [85]. It was reported that CAT protein levels and activity are elevated in the heart of mice that consumed a high-fat diet for only one day [85]. Thus, it appears that a robust switch to fatty acid oxidation is an adaptive cellular detoxification response to high-fat intake. It was proposed that since fatty acid oxidation enhances mitochondrial H_2_O_2_ production, high levels of CAT are needed to restore redox balance [85]. The findings of this study support the idea that obesogenic diets trigger an adaptive metabolic process in which CAT expression is upregulated to prevent cellular damage [86]. Furthermore, it indicates that catalase may represent a promising candidate for modulating redox dysregulation in obesity.

Glial cells have a high-energy demand, which is achieved by a strictly regulated cellular metabolism and high expression of redox proteins, including CAT [24,25,56]. The upregulation of GFAP in the rats exposed to the obesogenic WD and PS provides support to this idea. Several reports demonstrate that glial cells and CAT activities are particularly susceptible to the effects of high-fat diets and stress [20,22,87–89]. The brain redox state is sensitive to predator odor stress exposure in rats [73,74]. While these investigations show reduced CAT activities in the prefrontal cortex and amygdala of rats exposed to predator odor stress, the protein expression levels were not reported. It is possible that a reduction in CAT activity leads to increased CAT expression. Our data suggest that CAT upregulation may contribute to fear and anxiety-like phenotypes in rats exposed to an obesogenic diet and stress. This study supports the potential role of diet-induced redox dysregulation in behavioral correlates implicated in anxiety and stress-related disorders [90]. Oxidative stress could be a consequence of perturbations within several systems known to be disrupted in obesity and may be involved in predisposition and pathogenesis of stress-related disease.

### Limitations

The overarching hypothesis that abnormal fear network maturation and function in obesity implicates redox dysregulation/oxidative stress mechanisms should be further tested. Studies should be done to elucidate more specifically the molecular events that occur in WD-induced redox dysregulation, and more importantly, to establish causal relationships. Next investigations should also consider 1) using purified ingredient-matched control diets, 2) including female animals, and 3) testing the effects of DIO and PS on different rat strains. While this study did not provide redox status data from specific brain regions implicated in fear, whole-brain analyses provide fast and quantitative information on the general diet- and stress-induced redox status. Future studies are needed to validate the specific impact of obesogenic diets and stress on specific brain regions implicated with anxiety and stress-related disorders (e.g., hippocampus, prefrontal cortex, and amygdala).

### Conclusion

More comprehensive knowledge about the effects of obesity and poor dietary practices on stress reactivity can help to build up a more compelling panorama of the benefits of an early healthy antioxidant diet and, more importantly, help to identify biomarkers of psychiatric risk in affected obese individuals.

## Notes

### Competing Interest Statement

The authors have declared no competing interest.

